# Marine mammals and sea turtles listed under the U.S. Endangered Species Act are recovering

**DOI:** 10.1101/319921

**Authors:** Abel Valdivia, Shaye Wolf, Kieran Suckling

## Abstract

The U.S. Endangered Species Act (ESA) is the world’s strongest environmental law protecting imperiled plants and animals, and a growing number of marine species have been protected under this law as extinction risk in the oceans has increased. Marine mammals and sea turtles comprise 36% of the 161 ESA-listed marine species, yet analyses of recovery trends after listing are lacking. Here we gather the best available annual population estimates for all marine mammals (n=33) and sea turtles (n=29) listed under the ESA as species. Of these, we quantitatively analyze population trends, magnitude of population change, and recovery status for representative populations of 23 marine mammals and 9 sea turtles, which were listed for more than five years, occur in U.S. waters, and have data of sufficient quality and span of time for trend analyses. Using generalized linear and non-linear models, we found that 78% of marine mammals (n=18) and 78% of sea turtles (n=7) significantly increased after listing; 13% of marine mammals (n=3) and 22% of sea turtles (n=2) showed non-significant changes; while 9% of marine mammals (n=2), but no sea turtles declined after ESA protection. Overall, species with populations that increased in abundance were listed for 20 years or more (e.g., large whales, manatees, and sea turtles). Conservation measures triggered by ESA listing such as ending exploitation, tailored species management, and fishery regulations, among others, appear to have been largely successful in promoting species recovery, leading to the delisting of some species and to increases in most. These findings underscore the capacity of marine mammals and sea turtles to recover from substantial population declines when conservation actions under the ESA are implemented in a timely and effective manner.

## INTRODUCTION

Extinction risk for many marine species is increasing as the world’s ocean ecosystems are degraded by pervasive and increasing anthropogenic stressors [1,2] including over-fishing [3], habitat loss and degradation [4], pollution [5], and climate change [6,7]. Recent assessments have identified elevated levels of extinction risk in specific marine taxonomic groups: 14% of seagrasses [8], 16% of mangroves [9], 33% of reef-building corals [10], at least 25% of sharks and rays [11], and 11% of billfish and scombrids (e.g., tunas, bonitos, mackerels) [12]. Although considerably fewer extinctions of marine than terrestrial species have been recorded [1], marine species have a comparably high extinction risk as terrestrial species [13].

The Endangered Species Act (ESA) of the United States is the world’s strongest environmental law expressly designed to prevent extinction and promote recovery of imperiled species [14]. The strength of the ESA lies in its requirement to base decisions on the best available scientific information and its enforceable tools to reduce threats, protect habitat, and restore the abundance and geographic representation of listed species [15]. ESA’s tools include critical habitat designation, recovery planning with concrete and measurable goals, a science-based consultation process for federal agencies to prevent jeopardizing listed species or adversely modifying their critical habitat, and a prohibition on killing or harming listed species (16 U.S.C. § 1531 et seq.). Species protected under the ESA generally receive tailored federal and state conservation efforts with increased funding for management [16] and thus have better chances for recovery.

Evaluations of the ESA’s efficacy in preventing extinction and fostering recovery have become more imperative as extinction risks increase [1], available resources for conservation are often limited and mostly insufficient [17], and attacks on the ESA’s effectiveness by political opponents are escalating, with baseless critiques of the law [18]. Analyses to date of the ESA’s performance have consistently concluded that the ESA is highly effective in preventing species extinction [19]. After more than 45 years since the law was enacted in 1973, the ESA has shielded more than 99.5% of the species under its care from extinction [20]. Without the ESA’s protection, an estimated 227 species would have disappeared by 2006 [21].

The ultimate goal of the ESA is to promote the recovery of imperiled species. Numerous analyses have found that species status improves with time since listing, i.e., the longer a species is listed the more likely it is to increase in population abundance [22-24]. Species listed as threatened tend to respond faster to protection than endangered species because they generally have higher numbers at the time of listing, requiring relatively shorter time to recover [23,25]. Not surprisingly, species recovery is also associated with effectively implementation of the ESA’s tools, including funding for recovery actions [16,22,24,26,27]; presence of a dedicated recovery plan [23,28,29]; progress toward completing recovery goals (Abbitt & Scott 2001; Kerkvliet & Langpap 2007); and designation of critical habitat [23,but see 22,30,24].

Despite the increasing number of threatened and endangered marine species listed under the ESA [31], evaluations of the ESA’s track record in protecting marine species are lacking. This is especially evident for marine mammals and sea turtles that comprise a 36% of currently listed marine taxa [31]. Most studies of species recovery under the ESA are broad analyses of thousands of species [23,32-34] or are tailored to specific terrestrial-related taxa, such as plants [29], anadromous fish [35,36], amphibians [37], or birds [16,25,38,39]. Recent assessments of the status of marine mammal stocks in U.S. waters and global analyses of sea turtles categorize species by current population status, but do not analyze recovery trends since ESA listing [40,41].

The objective of our study was to assess how listed marine mammal and sea turtle species are faring under ESA protections, particularly for species occurring under U.S. jurisdiction where conservation actions promoted by the law are more robust. Thus, we gather the best available annual population estimates for all 33 marine mammals and 29 sea turtles listed under the ESA as species. Of these, we analyze recovery progress of 23 marine mammals and 9 sea turtles, which were listed for more than five years, reproduce or occur in U.S. waters, and had enough quality data to assess population trends during ESA protection. We hypothesize that marine mammals and sea turtles listed for several decades would be more likely to be recovering than recently listed species. To assess how ESA listing influenced population trends we followed three steps. First, we selected one representative population for each marine mammal and sea turtle species. Second, using generalized linear and non-linear models, we calculated species-specific population-level trends (significantly increased, no significant change, or significantly decreased) and magnitude of population change for each species after ESA protection. Third, we discuss conservation actions promoted by ESA listing that may contribute to species recovery, and illustrate this through three case-study species: the humpback whale in Hawaii and Alaska, Western Steller sea lion, and the North Atlantic green sea turtle.

## METHODS

### ESA listed marine mammal and sea turtle species selection

We reviewed the NMFS and USFWS’s endangered and threatened species database and selected all marine mammals (n=33) and sea turtles (n=29) currently listed or delisted as “species” under the ESA (Table 1, and S1 Table). We followed the ESA’s definition of “species”, which includes both species and subspecies that “interbreed when mature,” as well as distinct population segments (DPSs) (16 U.S.C. § 1532(16)). A DPS is defined as a vertebrate fish or wildlife population or a group of populations that is discrete from other populations of the species and is considered significant in relation to the entire species (61 FR 4722). For example, the humpback whale (*Megaptera novaeangliae*) is a biologically defined species, but it is currently divided in 14 DPSs under the ESA (81 FR 62259) that are considered different ESA-listed species.

**Table 1.**
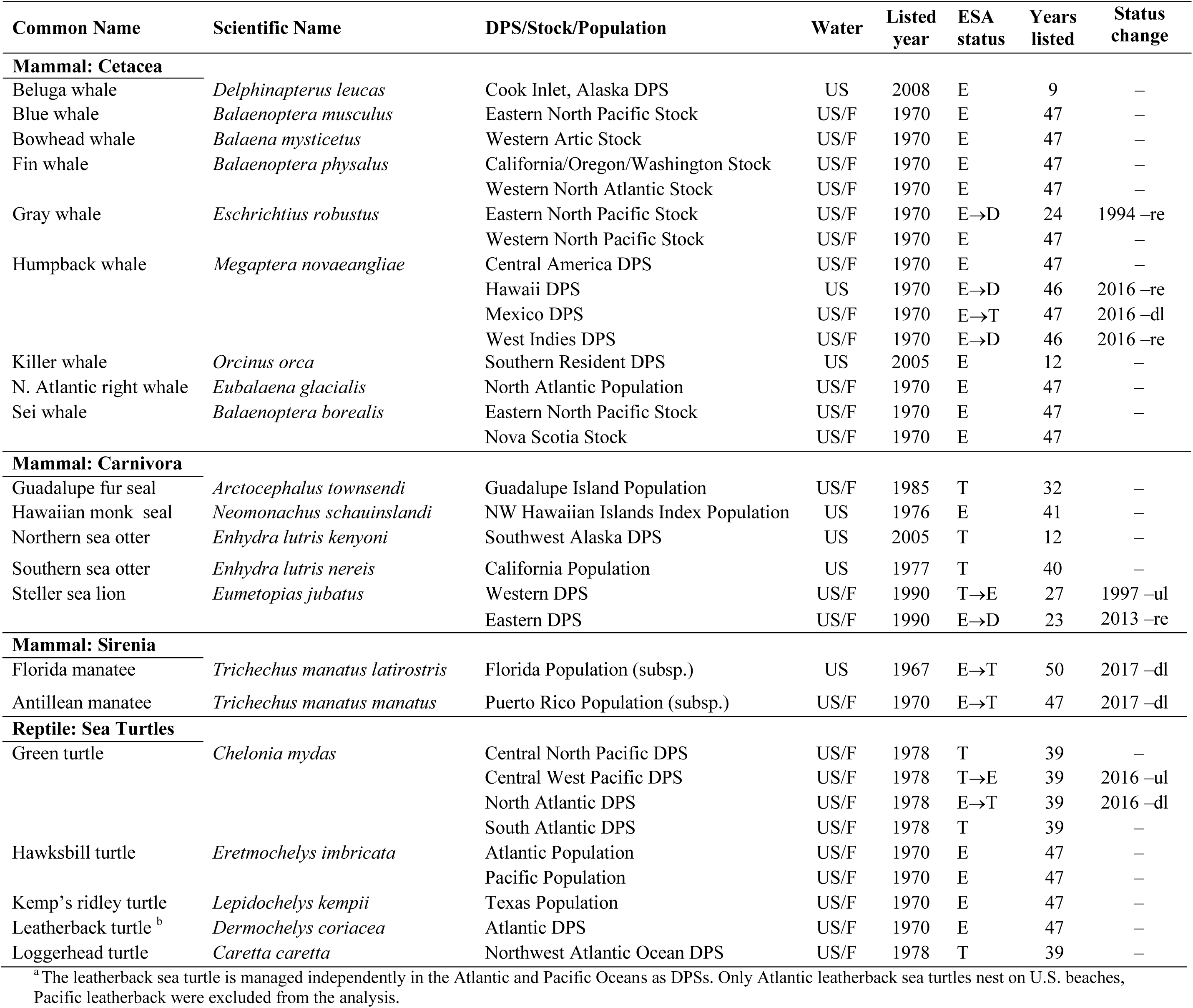
Status of marine mammals and sea turtle species protected under the ESA included in the analysis. These species are listed for more than five years (before 2012), are found exclusively within United States (US) or within both US and foreign (US/F) waters, have adequate population data that cover at least 40% of the listing period, and the population represents over 50% of the listed species. Distinct population segment (DPS); listing year; ESA status as endangered (E), threatened (T), delisted (D), or status change (e.g., T→E); and number of years listed are shown. Year of ESA status change due to down-listing (dl) and up-listing (ul); and reason for delisting such as recovered (re) are presented. Several species were listed before 1973 under the Endangered Species Preservation Act of 1966 and the Endangered Species Conservation Act of 1969, which were later replaced by the more comprehensive Endangered Species Act of 1973. See S1 Table for marine mammals and sea turtle ESA species excluded from the analyses. Data as of July 2017 [31].

| Common Name | Scientific Name | DPS/Stock/Population | Water | Listed year | ESA status | Years listed | Status change |
| --- | --- | --- | --- | --- | --- | --- | --- |
| **Mammal: Cetacea** |  |  |  |  |  |  |  |
| Beluga whale | *Delphinapterus leucas* | Cook Inlet, Alaska DPS | US | 2008 | E | 9 | – |
| Blue whale | *Balaenoptera musculus* | Eastern North Pacific Stock | US/F | 1970 | E | 47 | – |
| Bowhead whale | *Balaena mysticetus* | Western Artic Stock | US/F | 1970 | E | 47 | – |
| Fin whale | *Balaenoptera physalus* | California/Oregon/Washington Stock | US/F | 1970 | E | 47 | – |
|  |  | Western North Atlantic Stock | US/F | 1970 | E | 47 | – |
| Gray whale | *Eschrichtius robustus* | Eastern North Pacific Stock | US/F | 1970 | E→D | 24 | 1994 –re |
|  |  | Western North Pacific Stock | US/F | 1970 | E | 47 | – |
| Humpback whale | *Megaptera novaeangliae* | Central America DPS | US/F | 1970 | E | 47 | – |
|  |  | Hawaii DPS | US | 1970 | E→D | 46 | 2016 –re |
|  |  | Mexico DPS | US/F | 1970 | E→T | 47 | 2016 –dl |
|  |  | West Indies DPS | US/F | 1970 | E→D | 46 | 2016 –re |
| Killer whale | *Orcinus orca* | Southern Resident DPS | US | 2005 | E | 12 | – |
| N. Atlantic right whale | *Eubalaena glacialis* | North Atlantic Population | US/F | 1970 | E | 47 | – |
| Sei whale | *Balaenoptera borealis* | Eastern North Pacific Stock | US/F | 1970 | E | 47 | – |
|  |  | Nova Scotia Stock | US/F | 1970 | E | 47 |  |
| **Mammal: Carnivora** |  |  |  |  |  |  |  |
| Guadalupe fur seal | *Arctocephalus townsendi* | Guadalupe Island Population | US/F | 1985 | T | 32 | – |
| Hawaiian monk seal | *Neomonachus schauinslandi* | NW Hawaiian Islands Index Population | US | 1976 | E | 41 | – |
| Northern sea otter | *Enhydra lutris kenyoni* | Southwest Alaska DPS | US | 2005 | T | 12 | – |
| Southern sea otter | *Enhydra lutris nereis* | California Population | US | 1977 | T | 40 | – |
| Steller sea lion | *Eumetopias jubatus* | Western DPS | US/F | 1990 | T→E | 27 | 1997 –ul |
|  |  | Eastern DPS | US/F | 1990 | E→D | 23 | 2013 –re |
| **Mammal: Sirenia** |  |  |  |  |  |  |  |
| Florida manatee | *Trichechus manatus latirostris* | Florida Population (subsp.) | US | 1967 | E→T | 50 | 2017 –dl |
| Antillean manatee | *Trichechus manatus manatus* | Puerto Rico Population (subsp.) | US/F | 1970 | E→T | 47 | 2017 –dl |
| **Reptile: Sea Turtles** |  |  |  |  |  |  |  |
| Green turtle | *Chelonia mydas* | Central North Pacific DPS | US/F | 1978 | T | 39 | – |
|  |  | Central West Pacific DPS | US/F | 1978 | T→E | 39 | 2016 –ul |
|  |  | North Atlantic DPS | US/F | 1978 | E→T | 39 | 2016 –dl |
|  |  | South Atlantic DPS | US/F | 1978 | T | 39 | – |
| Hawksbill turtle | *Eretmochelys imbricata* | Atlantic Population | US/F | 1970 | E | 47 | – |
|  |  | Pacific Population | US/F | 1970 | E | 47 | – |
| Kemp's ridley turtle | *Lepidochelys kempii* | Texas Population | US/F | 1970 | E | 47 | – |
| Leatherback turtle <sup>b</sup> | *Dermochelys coriacea* | Atlantic DPS | US/F | 1970 | E | 47 | – |
| Loggerhead turtle | *Caretta caretta* | Northwest Atlantic Ocean DPS | US/F | 1978 | T | 39 | – |

<sup>a</sup>The leatherback sea turtle is managed independently in the Atlantic and Pacific Oceans as DPSs. Only Atlantic leatherback sea turtles nest on U.S. beaches,
Pacific leatherback were excluded from the analysis.

To determine the influence of ESA conservation measures on species recovery, we selected extant marine mammal and sea turtle species, listed or delisted, that meet five criteria: (1) ESA-listed for more than five years (pre-2012) to provide a minimum of post-listing population data time for conservation measures to be applied; (2) occurrence and reproduction in U.S. waters, i.e., excluding species that only occur in foreign waters where the ESA provides fewer protections [42]; (3) with enough reliable abundance data to determine population-level trends, i.e., at least three data points within 10 years, which is generally recommended for determining population change in ESA endangered and threatened species (82 FR 24944) and has been used for marine mammals [43] and sea turtles [44]; (4) with population data covering at least 40% of the ESA listing period, which we considered adequate for determining population trends after ESA listing; and (5) with a population that numerically represents over 50% of the abundance of the listed species (e.g., most green sea turtles of the North Atlantic DPS nest in Florida and thus Florida nest counts were used to represent this DPS). To delimit a population in our study, we used abundance data consistently collected over time in geographically defined areas such as stocks under the Marine Mammal Protection Act (MMPA), recovery units and DPSs as managed under the ESA, and other geographically restricted population units (e.g., ocean basins). As a result, population trend calculations are likely representative of the status of the entire ESA-listed species, even though an ESA-listed species may be comprised of several smaller populations. We identified 32 species that met our selection criteria, totaling 23 marine mammals and nine sea turtles (Tables 1 and S1). Of the 26 marine mammal and sea turtle species that did not meet our selection criteria, most (74%) do not occur in U.S. waters.

We also evaluated changes in species protection status. Species can be listed under the ESA as endangered or threatened. The ESA defines an endangered species as “in danger of extinction throughout all or significant portion of its range” while threatened species are “likely to become endangered in the foreseeable future throughout all or significant portion of its range” (16 U.S.C. § 1532(6) and (20)). For several species, the protection status (i.e., endangered or threatened) changed since the species was first listed at the global population level, and a few species were divided into DPSs (Table 1 and S1 Table). For the purpose of our study, we used the most current ESA protection status but the original year that the species was protected (Table 1).

### Data compilation and availability

We collected information and population-level abundance estimates for ESA-listed marine mammals and sea turtles from published papers and government reports. Main data sources included NMFS and USFWS technical memorandum and administrative reports, U.S. marine mammal stock assessment reports, species recovery plans, five-year status reviews, and primary sources from peer-reviewed scientific journals (see data in supporting information). When possible, we collected abundance data up to 2017 or to the most recently available population-level estimate. For populations that occur and reproduce in both U.S. and foreign waters, we used datasets from surveys that occurred, at least in part, in waters under U.S. jurisdiction.

Population abundance estimates came from a variety of survey methodologies (aerial, land, and ship-based surveys), mark-recapture population modeling, extrapolated data based on sex ratios, and photo-identification models (see data in supporting information). For marine mammals, population abundance comprised the total number of individuals including adults, juveniles, and pups or calves. For sea turtles, we used number of nests on nesting beaches, number of nesting females, or number of individuals to determine population trends. The number of nesting females and number of nests are common metrics for monitoring and evaluating population status of sea turtles [44].

Estimate bias and errors in population abundance obtained from data sources were variable among species and even within the same species over time. For example, survey effort and methodologies changed over time and population estimates have been calculated using different approaches over the years for the same population (e.g., traditional population abundance models, Bayesian population models, or habitat-based density models). Thus, where available, each data point was accompanied with information on data collection methodology, error information (e.g., coefficient of variation), and data estimation reliability (see data in supporting information). Time-series of population abundance for each species were carefully constructed to ensure all annual data points were derived from adequate and quantitative methodologies.

### Population trends and magnitude of change

For each marine mammal and sea turtle species, we calculated the population trend (as percentage change per year) and the magnitude of population change (as percentage change) after ESA listing based on the predicted distributions from the best and final fitted generalized linear and non-linear models (Table 2 and S2 Table). Population trajectories were classified as significantly increasing, non-significant change (non-significant slope), or significantly decreasing as in Magera et al. [43]. Recovering populations were defined as those that significantly increased in abundance after ESA listing, independently of whether or not they were on track to meet the recovery criteria for downlisting or delisting found in recovery plans. Populations with non-significant trends were not classified as “stable” as in other studies [see 40]. This was because determining population stability overtime requires further assessment of the accuracy of annual population estimates (e.g., the confidence intervals), which were often not available. Analysis of the magnitude of population recovery from estimated historical baselines was also not performed because this has been described elsewhere [43,45,46].

**Table 2.** Linear model and ANOVA results of the relationship between time since ESA listing and population trends (increasing, non-significant, decreasing) for marine mammal and sea turtle species. The decreasing trend was used as reference for the linear model. Significant codes are ‘**’ 0.01 and ‘*’ 0.05.

| **Linear model** | **Estimate** | **Std. Error** | **t value** | **Pr(>\|t\|)** |
| --- | --- | --- | --- | --- |
| (Intercept) | 26.500 | 7.327 | 3.617 | 0.00112 ** |
| Non-significant trend | 2.700 | 8.669 | 0.311 | 0.75769 |
| Increasing trend | 16.020 | 7.614 | 2.104 | 0.04417 * |
| **ANOVA** | **DF** | **Mean Sq** | **F value** | **Pr(>F)** |
| Trend | 2 | 548.17 | 5.1057 | 0.01260 * |
| Residuals | 29 | 107.36 |  |  |

### Data analysis: Population trajectories and model selection

To explore population trajectories after listing we used several models including linear models (*lm*), generalized linear models (*glm*), generalized least square models (*gls*), or generalized additive models (*gam*) in which population abundance estimates were modeled by running time in years (S2 Table). Because population trends were species specific, we used several family distributions and error links for each of the population-level models (S2 Table). For each population, we performed a comprehensive exploratory analysis using all models and possible combinations of families and links with and without a log transformation of the abundance estimates. In several *gls* models we added correlation and variance structures to account for potential temporal autocorrelation among years and variation in the data (S2 Table). Improvement in model fit was evaluated through theoretical model inference based on Akaike’s Information Criterion (AIC) [47], and comparing adjusted regression (r-squared) parameters when available [48]. Final model selection was based on a multi-model inference approach using AICc corrected for small samples [49]. See supporting information for final model details (S2 Table). All calculations and graphing were performed in R version 3.3 [50] using the packages *nlme v.3.1-131* for generalized least squared models [51]; *gam v.1.14-4* for generalized additive models [52]; *MuMIn v.1.15.6* for multi-model inference [53]; and *ggplot2 v.2.2.1* for data visualizations [54]. The dataset with specific data sources and references, and the R code of the analysis are provided in supporting information.

## RESULTS

### Status of ESA-listed marine mammal and sea turtle species in U.S. waters

Protection status for 10 out of the 32 species analyzed in our study changed since they were first listed, with eight of the 10 improving in status. Four species were downlisted from endangered to threatened: the Mexico DPS of humpback whale in 2016; the Florida manatee (*Trichechus manatus latirostris*) and the Antillean manatee (*Trichechus manatus manatus*) subspecies in 2017; and the North Atlantic DPS of green sea turtle in 2016 (Table 1). Four species were delisted from the ESA because NMFS determined they have recovered: the Eastern North Pacific Stock of gray whale (*Eschrichtius robustus*) in 1994, two DPSs of humpback whales (Hawaii and West Indies) in 2016, and the Eastern Pacific DPS of Steller sea lion in 2013 (Table 1). Two species were uplisted from threatened to endangered: the Western Pacific DPS of Steller sea lion (*Eumetopias jubatus*) in 1997, and the Central West Pacific DPS of green sea turtle in 2016 (Table 1).

### Population trends and magnitude of change

Overall, approximately 78% of marine mammals (18 out of 23) and 78% of sea turtles (7 out of 9) analyzed that met our selection criteria significantly increased in abundance after ESA listing (Fig 1A). Representative populations of three marine mammals (~13%) and two sea turtles (~22%) experienced non-significant change. Only two marine mammals (~9%), but no sea turtles significantly declined after ESA protection (Fig 1A). Marine mammals and sea turtles with populations that significantly increased were listed between two to five decades and increasing population trends was positively associated with time since listing. In contrast, there was no association with listing time for populations that showed non-significant trend or that declined in abundance (Fig 1B; Table 2). Out of the 25 species with populations that significantly increased, 52% were listed as endangered, 32% as threatened, and 16% were delisted, indicating that population increases occurred independent of whether a species was classified as threatened or endangered (Tables 1 and 2).

**Fig 1.**
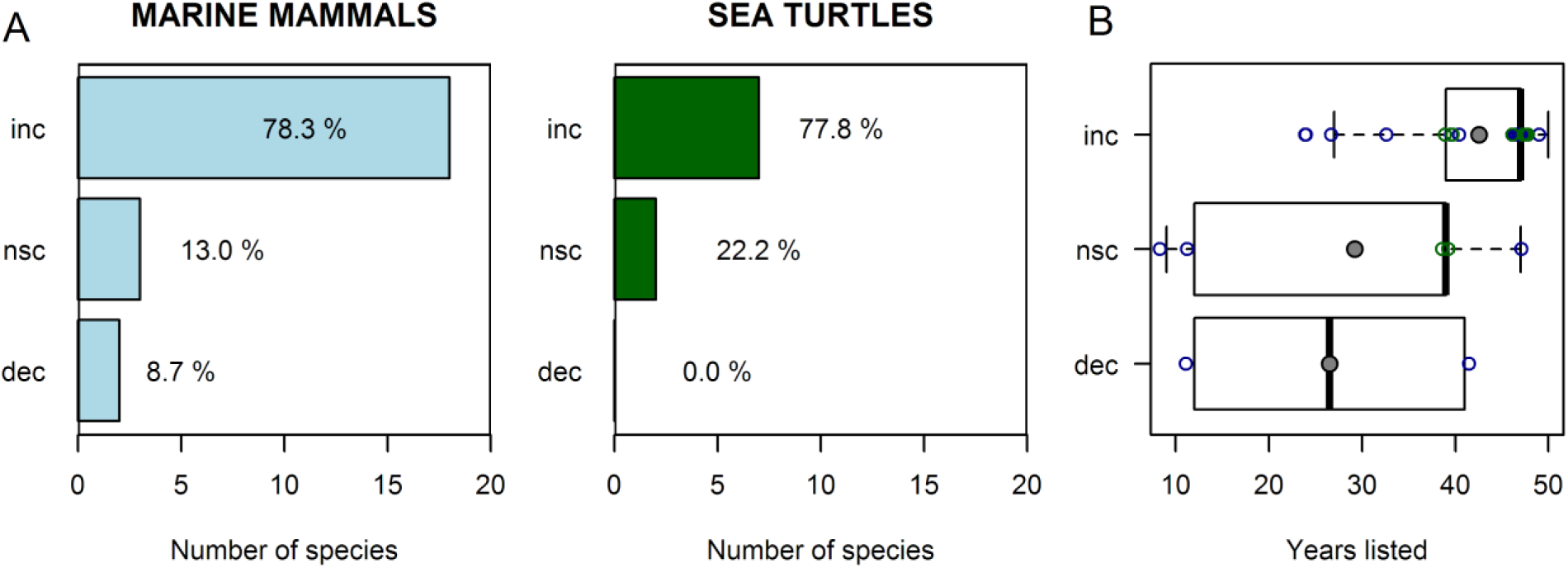
Number and percentage of marine mammal and sea turtle species protected under the ESA with population trends that significantly increased (inc), non-significant change (nsc), and significantly decreased (dec) after listing. **(A)** Calculations were based on representative populations of 23 marine mammal and 9 sea turtle ESA listed species that met our selection criteria. **(B)** Relationship between population trend and time since listing for marine mammals (blue circles) and sea turtles (green circles) species. Black line is the median and grey circle the mean.

Most marine mammals that significantly increased after ESA listing had substantial population growth (Figs. 2 and 3; Table 3). Several populations of large whale species increased in numbers from ~3% to ~43% per year, often doubling to quadrupling their initial population estimates (Table 3). For example, all four DPSs of humpback whales analyzed in our study showed substantial population increases (Fig 2; Table 3). In fact, the Hawaiian DPS of humpback whale reached over 10,100 individuals in 2005 from only 800 individuals estimated in 1979 (Fig 2; Table 3). NMFS subsequently delisted it from the ESA in 2016 (Table 1). While most large whale populations trended toward recovery, the critically endangered population of the North Atlantic right whale (*Eubalaena glacialis*) increased at 4.2% per year from 270 to 481 whales between 1990 and 2010, but declined to an estimated 451 whales between 2010 to 2016 (Fig 2; Table 3 and S2 Table). At least 17 individuals died in 2017, representing nearly 4% of the entire population (Table 3 and S2 Table).

**Fig 2.**
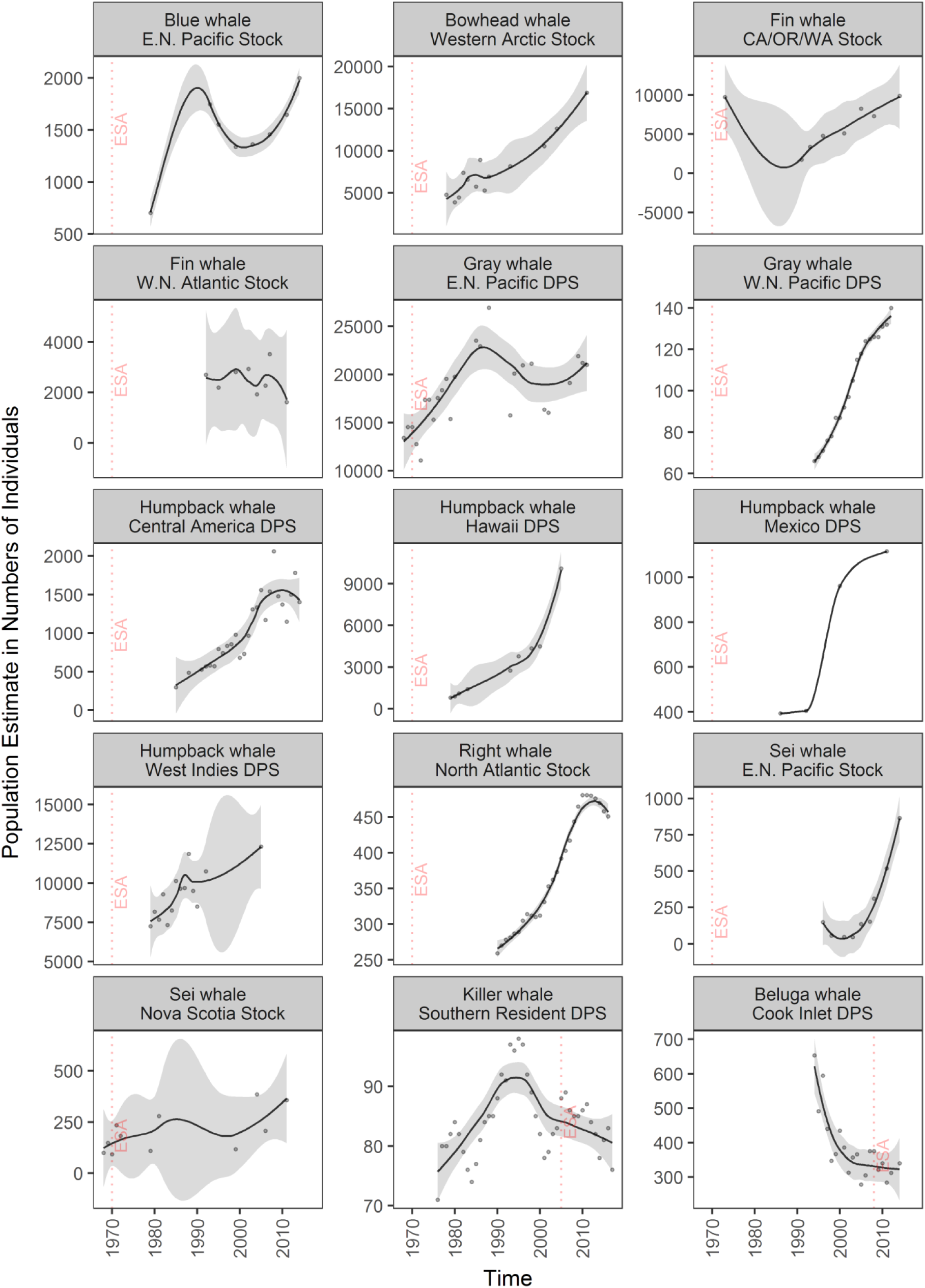
Population trends of cetacean marine mammals listed under the ESA. Trend lines (gray area: 95% confidence interval) are loess curves with span of 0.5 to aid in visual representation. Grey dots are estimated number of individuals. Panels are organized by decreasing length of time listed and then in alphabetical order based on species names. Dashed vertical red lines indicate the year of ESA listing. For species selection criteria see methods; for protection status see Table 1; and for results of fitting models see S2 Table. Abbreviations are CA/OR/WA: California/Oregon/Washington; E.N.: Eastern North; and W.N.: Western North.

**Fig 3.**
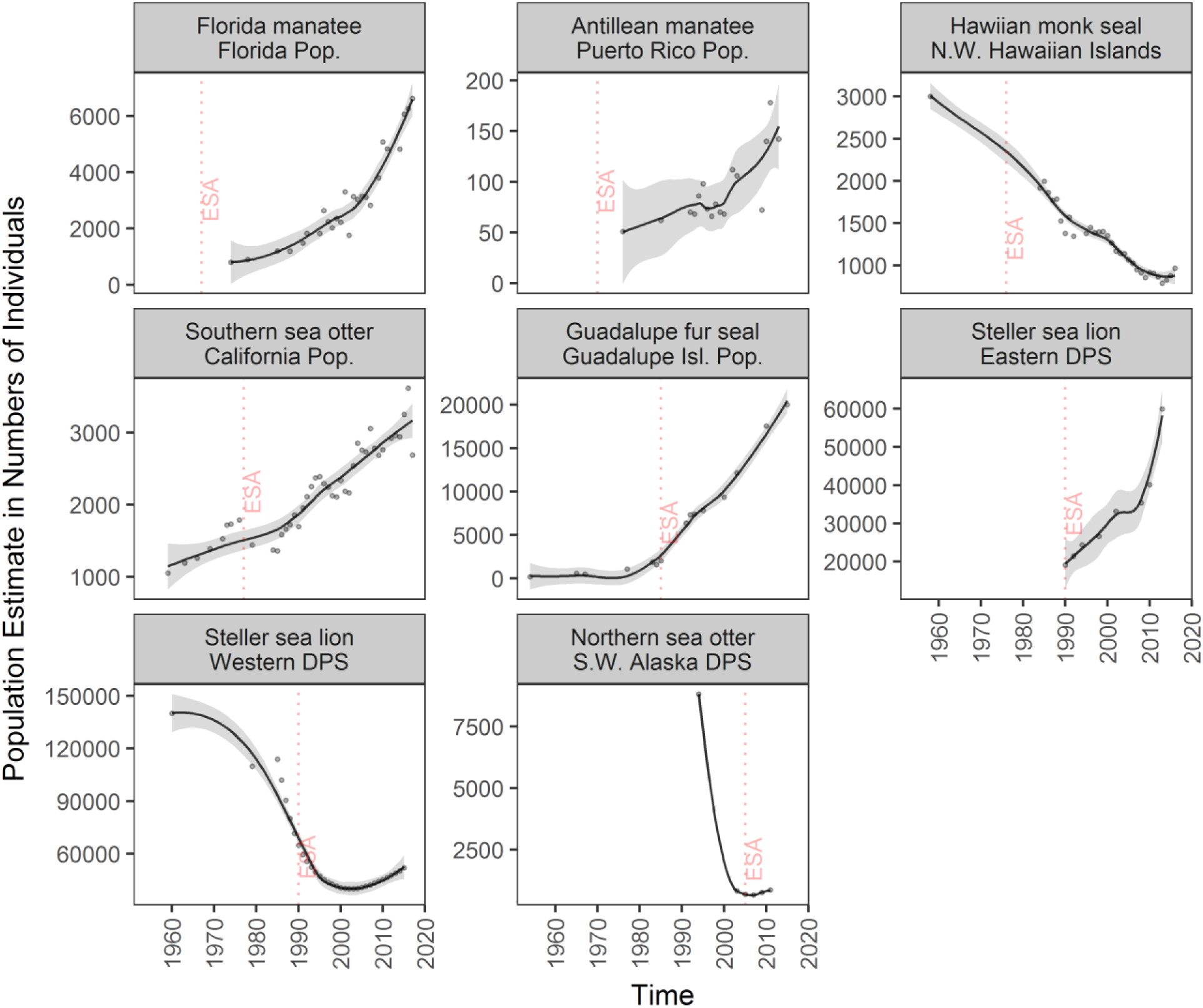
Population trends of non-cetacean marine mammals listed under the ESA. Trend lines (gray area: 95% confidence interval) are loess curves with span of 0.5 to aid in visual representation. Grey dots are estimated number of individuals. Panels are organized by decreasing length of time listed. Dashed vertical red lines indicate the year of ESA listing. For species selection criteria see methods; for protection status see Table 1; and for results of fitting models see S2 Table. Abbreviations are DPS: Distinct Population Segment; Pop.: Population; N.W. North Western; and S.W: Southwest.

**Table 3.** Population trends and magnitude of population change of selected marine mammal and sea turtle species protected under the ESA. Population (Pop.) trends (significantly increased↑, non-significant change →, significantly decreased↓) are based on species-specific models and time periods are shown. Current population trends (% per year) and magnitude of population change (%) were calculated based on available data after listing. First and last population abundance estimates for the time period are shown for reference.

| ESA Species (DPS/Stock/Location) | Time period (years) | Pop. trend (sign) | Pop. trend (% yr<sup>-1</sup>) | Pop. change (%) | First pop. estimate (No.) | Last pop. estimate (No.) |
| --- | --- | --- | --- | --- | --- | --- |
| **Cetacean** |  |  |  |  |  |  |
| Beluga whale (Cook Inlet DPS) | 08-14 | → | - 0.44 | - 8.8 | 375 | 340 |
| Blue whale (Eastern North Pacific Stock) | 79-14 | ↑ | + 4.99 | + 174.5 | 705 | 2,000 |
| Bowhead whale (Western Arctic Stock) | 78-11 | ↑ | + 8.34 | + 273.1 | 4,765 | 16,892 |
| Fin whale (California, Oregon, Washington Stock) | 91-14 | ↑ | + 13.34 | + 306.9 | 1,744 | 9,892 |
| Fin whale (Western North Atlantic Stock) | 92-11 | → | - 0.75 | -14.2 | 2,700 | 1,618 |
| Gray whale (Eastern North Pacific Stock) | 70-11 | ↑ | + 1.28 | + 52.6 | 14,553 | 20,990 |
| Gray whale (Western North Pacific Stock) | 94-12 | ↑ | + 6.22 | + 111.9 | 66 | 140 |
| Humpback whale (Central America DPS, California + Oregon) | 85-14 | ↑ | + 15.18 | + 440.2 | 300 | 1,403 |
| Humpback whale (Hawaii DPS, Hawaii winter) | 79-05 | ↑ | + 42.86 | + 1,114.3 | 800 | 10,103 |
| Humpback whale (Mexico DPS, Southeast Alaska to Alaska Peninsula) | 86-11 | ↑ | + 13.40 | + 334.4 | 393 | 1,115 |
| Humpback whale (West Indies DPS) | 79-05 | ↑ | + 3.00 | + 78.0 | 7,260 | 12,312 |
| Killer whale (Southern Resident DPS) | 05-17 | ↓ | - 0.93 | - 11.2 | 88 | 76 |
| North Atlantic right whale (North Atlantic) | 90-10 | ↑ | + 4.20 | + 84.0 | 270 | 481 |
|  | 10-16 | ↓ | - 1.05 | - 6.3 | 481 | 451 |
| Sei whale (Eastern North Pacific Stock) | 96-14 | ↑ | + 33.09 | + 595.6 | 150 | 864 |
| Sei whale (Nova Scotia Stock) | 70-11 | ↑ | + 1.98 | + 81.4 | 93 | 357 |
| **Carnivora** |  |  |  |  |  |  |
| Guadalupe fur seal (Guadalupe Island, Mexico) | 85-15 | ↑ | + 14.84 | + 905.4 | 2,017 | 20,000 |
| Hawaiian monk seal (NW Hawaiian Islands) | 85-13 | ↓ | - 2.04 | - 57.0 | 1,997 | 789 |
| (NW Hawaiian Islands) | 13-16 | ↑ | + 5.72 | + 22.9 | 789 | 968 |
| Northern sea otter (Southwest Alaska DPS,<br>Attu, Amchitka, Adak, Kiska Islands) | 05-11 | → | + 5.06 | + 30.3 | 687 | 863 |
| Southern sea otter (California) | 79-17 | ↑ | + 3.02 | + 114.7 | 1,443 | 2,688 |
| Steller sea lion (Eastern DPS,<br>California to Southeast Alaska) | 90-13 | ↑ | + 5.79 | + 133.2 | 19,103 | 59,968 |
| Steller sea lion (Western DPS, Alaska ) | 90-03 | ↓ | - 3.04 | - 39.4 | 64,761 | 39,963 |
|  | 03-15 | ↑ | + 2.34 | + 28.1 | 39,963 | 52,009 |
| **Sirenia** |  |  |  |  |  |  |
| Florida manatee (Florida) | 74-17 | ↑ | + 17.14 | + 737.3 | 800 | 6,620 |
| Antillean manatee (Puerto Rico) | 76-13 | ↑ | + 4.75 | + 175.8 | 51 | 142 |
| **Sea Turtles** |  |  |  |  |  |  |
| Green turtle (Central North Pacific DPS,<br>East Island, French Frigate Shoals, HI)<sup>1</sup> | 78-16 | ↑ | + 12.66 | + 480.9 | 101 | 88 |
| Green turtle (Central West Pacific DPS,<br>Guam waters)<sup>2</sup> | 78-10 | → | + 7.46 | + 238.6 | 92 | 299 |
| Green turtle (North Atlantic DPS,<br>Florida index beaches)<sup>3</sup> | 89-16 | ↑ | + 75.71 | + 2,044.2 | 464 | 2,978 |
| Green turtle (South Atlantic DPS,<br>Buck Reef NWR + Sandy Point NWR + Jack,<br>Isaac, and East End Bays, VI)<sup>3</sup> | 82-15 | ↑ | + 104.2 | + 3,439.1 | 31 | 931 |
| Hawksbill turtle (Atlantic DPS,<br>Mona Island, Puerto Rico)<sup>3</sup> | 74-15 | ↑ | + 22.64 | + 928.5 | 177 | 1,328 |
| Hawksbill turtle (Pacific DPS,<br>Island of Hawaii)<sup>1</sup> | 91-15 | ↑ | + 13.91 | + 333.9 | 4 | 25 |
| Kemp's ridley turtle (Texas)<sup>3</sup> | 79-17 | ↑ | + 284.2 | + 11,083.8 | 1 | 353 |
| Leatherback turtle (Atlantic DPS,<br>Florida + Puerto Rico + Sandy Point NWR, VI)<sup>3</sup> | 84-16 | ↑ | + 32.25 | + 1,032.2 | 368 | 3,625 |
| Loggerhead turtle (NW Atlantic DPS,<br>Peninsular FL index beaches)<sup>3</sup> | 89-16 | → | + 1.16 | + 31.4 | 39,083 | 65,807 |

<sup>1</sup> Number of nesting females
<sup>2</sup> Number of individuals
<sup>3</sup> Number of nests

Non-cetacean marine mammals also significantly increased in abundance at relatively high population growth rates since ESA protection. Notably, the Guadalupe fur seal (*Arctocephalus townsendi*) increased about nine times at ~15% per year since listing in 1985 (Fig 3; Table 3). The California population of the Southern sea otter (*Enhydra lutris nereis*) approximately doubled in numbers and it is likely to reach the demographic recovery criteria in the coming years (Fig 3; Table 3). The Eastern DPS of Steller sea lion (*Eumetopias jubatus*) tripled its population at ~6% per year since 1990, reaching its recovery criteria of ~60,000 individuals in 2013, and was subsequently delisted from the ESA (Fig 3; Table 3). Also, both the Florida and Antillean manatee subspecies increased approximately eight and three times (~17% and ~5% per year), respectively, for the past 40 years (Fig 3; Table 3); and the USFWS downlisted them from endangered to threatened in 2017 (Table 1).

Representative populations of five marine mammals analyzed in our study did not increase in abundance. Three marine mammals showed non-significant population change after listing: Western North Atlantic fin whale (*Balaenoptera physalus*), Cook Inlet beluga whale (*Delphinapterus leucas*) DPS (Fig 2; Table 3 and S2 Table), and Southwest Alaska DPS of the northern sea otter (*Enhydra lutris kenyoni*) (Fig 3; Table 3 and S2 Table). In contrast, two marine mammals significantly declined after ESA listing: the critically endangered Southern Resident killer whale (*Orcinus orca*) and the Hawaiian monk seal (*Neomonachus schauinslandi*). Southern Resident killer whales declined at – 0.93% per year since listing in 2005, when the population had 88 individuals (Fig 2, Table 3). This population suffered major declines after a record high of 98 individuals in 1995, and the last population survey estimated 76 individuals in September 2017, a 30-year low (Fig 2; Table 3). Total abundance of Hawaiian monk seals from six index subpopulations in the Northwestern Hawaiian Islands significantly declined from 1,997 individuals in 1985 to 789 seals in 2013 at approximately – 2% per year (Fig 3; Table 3). However, the population has increased to 968 seals by 2016 (Table 3).

Nearly all sea turtle species analyzed in our study significantly increased after ESA listing (Fig 4; Table 3 and S2 Table). Estimates of the number of individuals, nesting females, and number of nests in nesting beaches of representative populations of green, hawksbill, Kemp’s ridley, and Atlantic leatherback sea turtles showed that these species increased at considerably high growth rates (~13% to ~284% per year) for several decades, depending on initial estimates (Fig 4; Table 3 and S2 Table). For example, the number of nesting females of green sea turtle at East Island of the French Frigate Shoals in Hawaii (Central North Pacific DPS) increased at ~13% per year from 101 individuals in 1978 to 492 nesting females in 2015 (Fig 4; Table 3). The number of nests of green sea turtles across Florida statewide beaches (North Atlantic DPS) increased at ~76% per year from 62 nests in 1979 to a record high of 37,341 nests in 2015 (Fig 4; Table 3). Due the strong recovery of green sea turtles across Florida, NMFS and the USFWS downlisted the entire North Atlantic DPS from endangered to threatened in 2016 (Table 1). Similarly, the number of nests of the hawksbill turtle population at Mona Island in Puerto Rico (Atlantic DPS) increased at over 22% per year from 177 in 1974 to a record high of 1,626 nests in 2014 (Fig 4; Table 3). Notably, the Atlantic leatherback populations have also experienced a considerable rebound, and the combined number of nests across Florida, Puerto Rico, and the Virgin Islands, significantly increased after ESA listing (Fig 4; Table 3).

**Figure 4.**
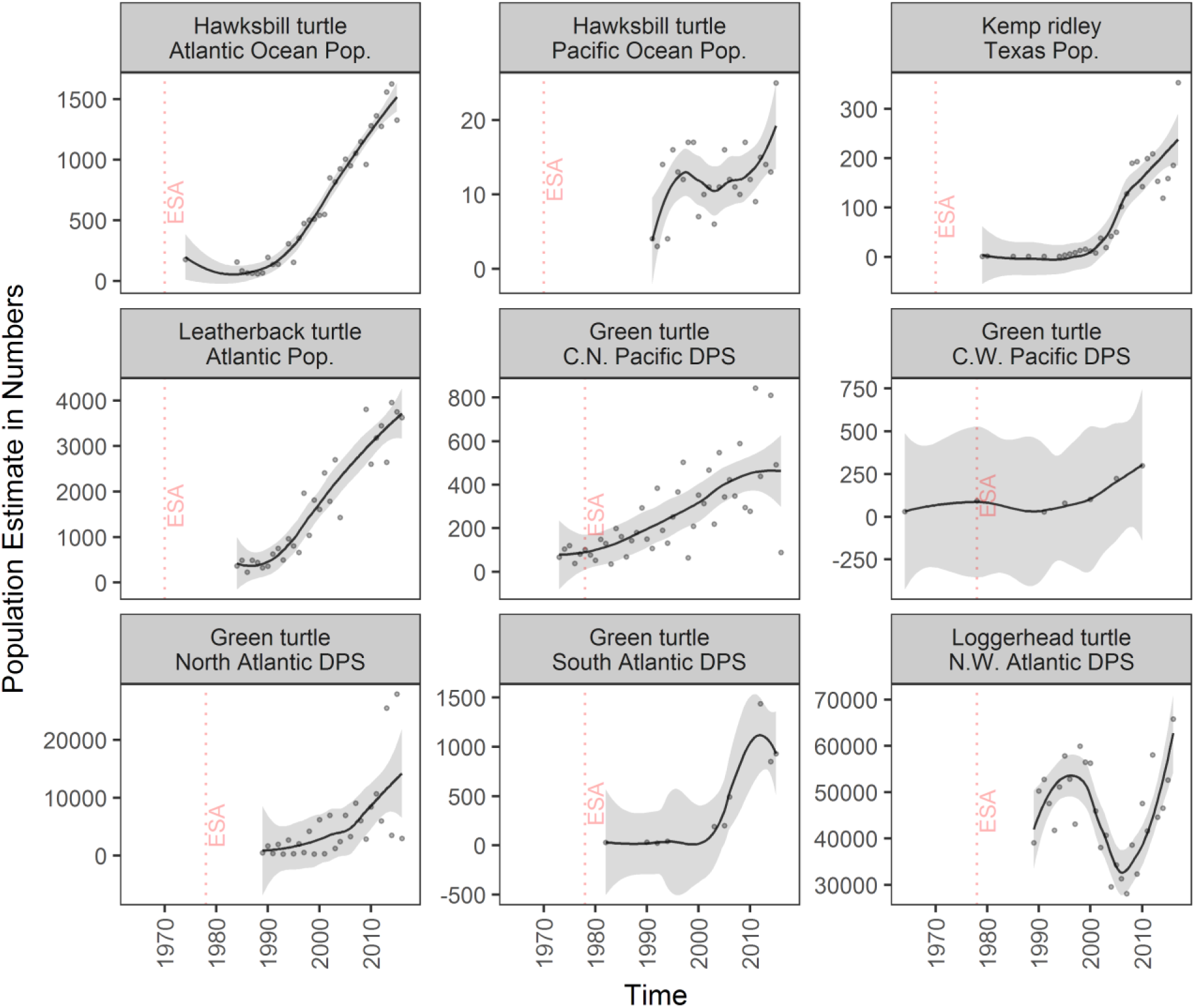
Population trajectories of sea turtles listed under the ESA. Trend lines (gray area: 95% confidence interval) are loess curves with span of 0.5 to aid in visual representation. Grey dots are estimated number of nests, except for the hawksbill Pacific DPS and the green turtle Central North Pacific DPS (number of nesting females), and green turtle Central West Pacific DPS (number of individuals). Panels are organized by decreasing length of time listed. Dashed vertical red lines indicate the year of ESA listing. For species selection criteria see methods; for protection status see Table 1; and for results of fitting models see S2 Table. Abbreviations are DPS: Distinct Population Segment; Pop.: Population; C.N.: Central North; C.W.: Central West; and N.W: Northwest.

Among the sea turtles analyzed in this study, our models were not able to detect significant population linear trends for the Central West Pacific DPS of the green turtle (Guam waters), and the Northwest Atlantic DPS of the loggerhead turtle (*Caretta caretta*) across the Florida peninsula (Fig 4; Table 3 and S2 Table). For example, the best models for the number of nests of loggerhead turtles across index beaches of the Florida peninsula described a non-linear relationship where the number of nests substantially fluctuated since 1989, with a record high of 65,807 in 2016 (Fig 4).

## DISCUSSION

Most marine mammals and sea turtles listed under the ESA that met our selection criteria, significantly increased after listing indicating strong population recoveries. Our analyses confirm the hypothesis that ESA listed marine mammal and sea turtle species are more likely to be recovering the longer they stay protected under the law, regardless of whether they are listed as threatened or endangered. Previous studies support these findings for a variety of terrestrial taxa, marine birds, and anadromous fishes [16,19,23,25,39,55]. Here we discuss how these findings suggest that ESA protections and conservation measures are important for the recovery of imperiled marine mammals and sea turtles, and illustrate specific examples through three case studies.

The ESA’s prohibitions on commercial exploitation paired with the implementation of widespread conservation measures such as interagency consultation, recovery plans and critical habitat designations have been crucial to mitigating threats that affect marine mammals and sea turtles [34,56]. Strong population increases for most marine mammal and sea turtle species after ESA protection demonstrate the capacity of these taxa to recover from drastic declines after decades of exploitation, habitat degradation, and other threats. Between the 18^th^ to early 20^th^ century these groups were substantially depleted [4,46,57,58], in a few cases to extinction such as the Steller’s sea cow [59] and the Caribbean monk seal [60,61]. Marine mammals and sea turtles have greatly benefited from a major change from resource exploitation (e.g., whaling, hunting, egg harvesting) to conservation measures that protect these species from direct and indirect harm [62].

For the large whales, ESA protections facilitated the recovery of populations that were severely depleted by commercial whaling by reducing key threats such as ship strikes, entanglement in fishing gear, and pollution [56,63-67]. For example, ESA protection led to the establishment of vessel speed limits and restrictions on approaching whales too closely to lower the likelihood of death and injury from vessel strikes [68-70]. By triggering a depleted designation under the U.S. Marine Mammal Protection Act (MMPA), ESA marine mammal listings have prompted the implementation of take reduction plans to reduce injury and death from fisheries entanglement that require gear modifications, time and area closures, and vessel observers [40,71]. ESA regulations have also helped to limit acoustic harms to whales and other marine mammals by restricting U.S. military use of sonar and explosions in biologically important habitat areas around Hawaii Islands and Southern California [72].

For sea turtles, ESA protections have been instrumental in reducing primary threats from human harvest, fishery bycatch, and habitat destruction. The ESA’s prohibitions on harvesting sea turtles and their eggs has virtually eliminated this key threat – historically the principal cause of sea turtle population declines – in U.S. turtle nesting and foraging grounds [73,74]. ESA listing prompted regulations that have reduced sea turtle bycatch mortality in commercial fisheries by requiring gear modifications (e.g., turtle excluder devices in trawl fisheries, circle hooks in longline fisheries, modifications to pound net leaders), time and area closures, bycatch limits, changes to fishing practices, and monitoring programs [75-78]. ESA-prompted reductions in off-road-vehicle use and night lighting on nesting beaches have improved nesting success [79,80], as has protection of important turtle nesting beaches as National Wildlife Refuges (NWR) on the Atlantic coast (e.g., Archie Carr NWR, Florida) and the U.S. Caribbean (e.g., Culebra NWR, Puerto Rico; Sandy Point NWR, U.S. Virgin Islands) [73,74,81]. In addition, ESA protections have facilitated federal and state agencies (e.g., National Park Service, Florida Fish and Wildlife Conservation Commission) to contribute funding and supported conservation efforts including species reintroductions, e.g., Kemp’s ridley turtles in Texas [82], and volunteer monitoring and scientific data collection on most sea turtle nesting beaches across the U.S. (e.g., Florida Statewide and Index Nesting Beach Survey program).

Two marine mammal species with populations that did not significantly change were listed relatively recently (< 15 years). For example, the Cook Inlet DPS of beluga whale in Alaska was listed in 2008 and the Southwest Alaska DPS of the northern sea otter was listed in 2005. Conservation measures for these two species were developed relatively recently and ongoing threats have not been mitigated [83,84]. It is likely that these species will require more time under ESA protection as well as the adoption of robust conservation measures. In contrast, one marine mammal and two sea turtles listed for several decades had populations with non-significant change. The lack of significant population changes in the Western North Atlantic Stock of fin whale and the Central West Pacific DPS of green turtle may be related to lack of statistical power to detect a trend in abundance as confidence intervals of population estimates were relatively large (Figs. 2 and 4; S1 Table) [74,85]. Alternatively, the populations of these species may be stable, but further population estimates are needed to determine stability [74,85]. Finally, fluctuations in the number of nests of the Northwest Atlantic DPS of loggerhead turtle across Florida beaches have been strongly correlated with ocean conditions associated with long term climate forcing such as the Atlantic Multidecadal Oscillation [86].

Endangered marine mammal species with relatively low population abundance that significantly declined after listing (e.g., Southern Resident killer whale and Hawaiian monk seal) or showed non-significant change (e.g., Cook Inlet beluga whale) require urgent conservation attention. NMFS already recognizes these species among those most at-risk of extinction in the immediate future and they are considered recovery priorities because of rapid population declines [87]. These species face several similar regional anthropogenic threats including prey reduction due to fishing, habitat degradation, toxic pollutants, disturbance from boat traffic and marine noise, fishery interactions, as well as global threats associated with climate change and ocean regime shifts that affect food availability [88-92]. In particular, food limitation has been recognized as a key driver of lower body condition, pregnancy failures, calf/pup and juvenile mortality, and lack of population recovery [92-96]. Numerous conservation measures addressing anthropogenic stressors have been developed for these species and are delineated in recovery plans [84,97,98]. For example, NMFS established regulations to protect killer whales in Washington waters from vessel impacts in 2011 (76 FR 20870). For Hawaiian monk seals, entanglements in fishing gear, fishery interactions, and other human-caused mortalities (e.g., intentional killing) have been reduced since ESA listing, especially across the inhabited Main Hawaiian Islands [98,99]. In fact, after more than 50 years of continued decline, the range-wide population seems to have steadily increased in numbers since 2013 to approximately 1,400 seals in 2016 [100]. Recently, stronger conservation measures have been developed in high-priority action plans that focus efforts and resources to reduce threats and stabilize population declines [87]. The outcomes of these conservation efforts will require time to be realized, although the compounding effects of climate change stressors may compromise the ability of these endangered species to rebound.

### Case studies illustrate the recovery benefits of ESA listing

#### Hawaii DPS of humpback whale

The Hawaii DPS of humpback whale was delisted by NMFS in 2016 based on its strong population growth and the mitigation of key threats (NMFS 2015). Whales in this population use the waters surrounding the main Hawaiian Islands for mating and calving and migrate to feeding in areas off Southeast Alaska and northern British Columbia. The size of the population in Hawaiian waters increased from 800 individuals in 1979 to more than 10,000 individuals in 2005, with the recent population growth rate estimated around 6% (NMFS 2015). ESA listing in 1970 prompted conservation measures in Hawaii and Alaska to reduce key threats to recovery. ESA regulations restricted vessels in Hawaiian and Alaskan waters from approaching whales within 100 yards, prohibited disrupting normal behaviors, and required slower vessel speeds to reduce the likelihood of ship strikes and minimize human disturbance (60 FR 3775, 66 FR 29502). ESA listing also prompted coordinated federal and state efforts to reduce whale entanglements in fishing gear through the Hawaiian Islands Disentanglement Network and Alaska Response Network. The threatened status of humpback whales also provided impetus for the designation of the 1,400 square-mile Hawaiian Islands Humpback Whale National Marine Sanctuary in 1992 to protect humpback whales and their habitat (60 FR 48000).

#### Western DPSs of Steller sea lion

Population abundance of the Western DPS of Steller sea lion, which ranges from Eastern Gulf of Alaska to the Western Aleutian Islands and Bering Sea [101], significantly increased over the past 13 years (Fig 3). This species has shown a tremendous population recovery despite years of overexploitation (for their fur, meat, and oil), indiscriminate killing, and decades of habitat degradation, ship strikes, and fishery interactions [102]. Subpopulations of the Western DPS suffered major declines through 2003 when a substantial population rebound began to occur [101]. Abundance estimates of the Western DPS declined from 140,000 to 110,000 individuals between 1960 and 1979 in rookeries and haul-outs across Southwest Alaska [102]. Total counts continued to decline at 15% per year in the late 1980s, prompting NMFS to list the Western DPS as threatened throughout the entire range in 1990 (NMFS 2008) and to uplist it to endangered in 1997 because of continued declines during the 1990s [103]. Population abundance stabilized in the early 2000s [104,105] with the lowest population estimate in 2003 [101]. Notably, population abundance significantly increased at 2.34% per year from 2003 to 2015 (Fig 5).

Conservation efforts under both the ESA and the MMPA such as designation of protective zones, critical habitat designation, fishery regulations for prey species, and local regulations around major rookeries and haul-outs have likely contributed to the species recovery success [102]. NMFS implemented several fishery management measures (e.g., area closures, catch and harvest limits, reduction of disturbance due to fishing) in the Alaska groundfish fisheries in 2003 (Bering Sea and Gulf of Alaska) around major haul-outs and rockeries within the designated critical habitat (68 FR 204). These regulations, designed to reduce competition for prey between commercial fishing and Steller sea lions and increase prey availability, are thought to have contributed to increased prey abundance and a rebound of the DPS [102,106]. In fact, counts have increased at an average of 2.17% (juveniles and adults) and 1.76% (pups) per year from 2000 to 2015 [101,107], although geographical variation exists due to migration among subpopulations (NMFS 2015).

#### North Atlantic DPS of green sea turtle

The North Atlantic DPS of the green sea turtle, which mostly nests across Florida beaches, is another ESA conservation success. The species has been increasing exponentially and has become one of the largest nesting aggregations in the western Atlantic in recent years [74]. Historically exploited across the Caribbean [46], this species has shown a high recovery potential when nesting areas are strictly protected from human disturbance and development, and fishery bycatch is substantially reduced [74]. The North Atlantic DPS of green turtles showed high records of nest numbers in 2013 (36,169 nests) and 2015 (37,341 nests) across Florida nesting beaches compared with only 62 nests estimated in 1979 (Fig 4). In 2016, NMFS and USFWS reclassified green sea turtles into 11 DPSs of which the Florida population was downlisted from endangered to threatened due to strong population growth and record numbers in nesting beaches throughout the peninsula (81 FR 20057).

ESA tools have been crucial for the recovery of the North Atlantic DPS of green sea turtles. ESA regulations have led to fishing gear modifications, major changes in fishing practices, time and area closures, and the establishment of turtle excluder devices for shrimp trawlers [77,108]. In particular, fishery regulations instituted because of ESA protection have been largely successful in reducing green sea turtle bycatch from Atlantic pelagic longlines and gillnets, the Chesapeake Bay pound net fishery, and the Gulf of Mexico’s shrimp and flounder trawl fisheries [74]. Take prohibitions (i.e., no killing of adults or egg harvesting) under the ESA have been a major conservation tool that directly boosts population growth. In addition, several national wildlife refuges were dedicated to protecting nesting areas on the Atlantic coast and Gulf of Mexico, with nest watchers and patrolling during nesting seasons [74]. The Florida Statewide Nesting Beach Survey program, initiated in 1979 (one year after listing) as a cooperative agreement between the USFWS and the Florida Fish and Wildlife Conservation Commission, now monitors ~215 nesting beaches (~825 miles) across Florida, involving federal, state, and regional institutions as well as several conservation organizations, university scientists, and private citizens [109]. Federal agencies (NMFS and USFWS) along with state agencies and other institutions have worked together in implementing the management actions in the 1991 recovery plan, eliminating or reducing threats in nesting and foraging areas [74].

## CONCLUSIONS

Recovery is occurring for most marine mammals and sea turtles listed under the ESA and analyzed in our study. Species listed for over 20 years were more likely to have populations that significantly increased in numbers. In contrast, relatively recently listed species were more likely to have populations with non-significant change or declining. Targeted conservation efforts triggered by ESA listing have been largely successful in promoting species recovery leading to the delisting of some species and to dramatic increases in most. The recovery of listed species depends ultimately on the adequate implementation of the ESA’s tools and conservation measures. Studies have demonstrated that the government’s failure to fully implement the ESA’s protections and adequately fund conservation actions have been major impediments to species recovery [19]. In general, listed species with designated critical habitat, sufficient conservation funding, and well-implemented species-specific recovery plans tend to recover relatively faster [16,23,24]. Our analysis not only underscores the capacity of marine mammals and sea turtles to rebound after decades of exploitation and habitat degradation, but also highlights the success of marine species conservation through the ESA protection.

## ACKNOWLEDGMENTS

Several staff from the Center for Biological Diversity gathered and organized the data used in this paper. Thanks to the thorough review by Miyoko Sakashita that substantially improved this paper.

